# Deep Learning Image Classification of Red Blood Cell Deformability

**DOI:** 10.1101/2021.07.26.453886

**Authors:** Erik S. Lamoureux, Emel Islamzada, Matthew V. J. Wiens, Kerryn Matthews, Simon P. Duffy, Hongshen Ma

**Author notes:** **Corresponding Author** Hongshen Ma, 2054-6250 Applied Science Lane, Vancouver, BC, Canada V6T 1Z4. **Data availability statement** The data that support the findings of this study are available on request from the corresponding author. **Ethics approval statement** This study was approved by the University of British Columbia’s Clinical Research Ethics Board (UBC REB# H19-01121) and Canadian Blood Services Research Ethics Board (CBS REB# 2019-029). **Author contributions** H.M. supervised the study. H.M., E.L., and S.P.D. conceived the idea. E.L., E.I., and K.M. performed the experimental work. E.L. and M.W. performed the computational work. All authors wrote the manuscript.

## Abstract

Red blood cells (RBCs) must be highly deformable to transit through the microvasculature to deliver oxygen to tissues. The loss of RBC deformability resulting from pathology, natural aging, or storage in blood bags can impede the proper function of these cells. A variety of methods have been developed to measure RBC deformability, but these methods require specialized equipment, long measurement time, and highly skilled personnel. To address this challenge, we investigated whether a machine learning approach could be applied to determine donor RBC deformability using single cell microscope images. We used the microfluidic ratchet device to sort RBCs based on deformability. Sorted cells are then imaged and used to train a deep learning model to classify RBCs based on deformability. This model correctly predicted deformability of individual RBCs with 84 ± 11% accuracy averaged across ten donors. Using this model to score the deformability of RBC samples were accurate to within 4.4 ± 2.5% of the value obtained using the microfluidic ratchet device. While machine learning methods are frequently developed to automate human image analysis, our study is remarkable in showing that deep learning of single cell microscopy images could be used to measure RBC deformability, a property not normally measurable by imaging. Measuring RBC deformability by imaging is also desirable because it can be performed rapidly using a standard microscopy system, potentially enabling RBC deformability studies to be performed as part of routine clinical assessments.

## INTRODUCTION

Red blood cells (RBCs) are highly specialized cells that facilitate tissue respiration by delivering oxygen and removing carbon dioxide. ^1,2^ RBCs transverse through the entire circulatory system approximately every 60 seconds. Their journey includes the microvasculature, where RBCs must deform through capillaries measuring as little as 2 µm in diameter, as well as the inter-endothelial clefts of the spleen measuring 0.5-1.0 µm in diameter. ^3,4^ The loss of RBC deformability, due to pathology, natural aging, or storage in blood bags, reduces the ability of RBCs to circulate and facilitate their removal from circulation by phagocytes in the spleen and the liver. ^5,6^ As a result, there is significant interest in methods for measuring RBC deformability as a potential biomarker for diseases, such as malaria ^7^ and hemoglobinopathies, ^2,8^ or for assessing the quality of donated RBCs for use in blood transfusions. ^9,10^

Approaches for measuring RBC deformability can be classified as either flow-based or deformation-based methods. Flow-based methods deform RBCs using fluid shear stress and then measure the resulting shape change. A classical method is ektacytometry, which deforms RBCs using shear flow between two transparent cylinders and then uses optical diffraction to measure the resulting population RBC elongation. ^11,12^ Other flow-based methods deform RBCs using high shear flow through microchannels and then measure the resulting RBC elongation using high speed imaging ^13,14^ or electrical impedance. ^15^ Classical deformation-based methods include micropipette aspiration, ^16^ atomic force microscopy, ^17^ and optical tweezers, ^18,19^ which measure RBC deformability based on partial or complete transit of each cell through microscale structures. Microfluidic deformation-based methods measure RBC deformability via capillary obstruction, ^20^ deposition length in tapered constrictions, ^21,22^ transit pressure through constrictions, ^10,23–28^ transit time through constrictions, ^29–31^ and sorting RBCs based on deformability using microfluidic ratchets. ^32–34^ A common challenge for all existing deformability assays is the needs for specialized apparatus and skilled personnel, which limit the ability to translate the technology to clinical settings. ^35^ Additionally, since different assays rely on different underlying principles for measuring RBC deformability, it is often difficult or impossible to compare results across studies.

As an alternative to physical measurement of RBC deformability, cues for biophysical changes in these cells may be extracted from cell imaging, without the need for highly specialized equipment and personnel. RBCs typically exhibit a highly deformable biconcave discoid morphology, and deviation from this morphology may correspond with changes in cell deformability. ^36,37^ In fact, deep learning methods have been developed to assess changes in RBC morphology during cold storage, ^38^ malaria, ^39–44^ sickle cell disease, ^45–50^ and thalassemia. ^51–53^ However, RBC morphology varies over the life cycle of the cell and this variability may obscure efforts to infer deformability from cell morphology. Furthermore, no specific morphological features can be directly attributed to predictable changes in RBC deformability. We recently developed a microfluidic process for deformability-based sorting of RBCs ^32–34^ as well as a deep learning method to distinguish cell lines based on feature differences imperceptible to human cognition. ^54^ We hypothesized that a combination of these advances could enable the direct measurement of RBC deformability by cell imaging using optical microscopy.

Here, we investigate the potential to use deep learning to estimate RBC deformability using brightfield microscopy images. We leverage the ability of a microfluidic ratchet device to sort RBCs based on deformability to generate training sets of RBCs with distinct deformability. We show that the deep learning model can classify RBC into *deformable* or *rigid* fractions using donor dataset sizes ranging from 20,000 to 70,000 images. For a sample of ten donors, who were diverse in terms of blood type and sex, testing classification accuracy ranged from 68-98% with an aggregate mean (± SD) of 84 ± 11%. Using our model to predict donor RBC Rigidity Scores (RS) was accurate to within a mean of 4.4 ± 2.5% deviation, compared to measurement using a microfluidic device. Our results confirm that RBC deformability can be estimated from microscopy images to potentially simplify the measurement of RBC deformability.

## RESULTS

### Approach

Our experimental approach (**Fig. 1**) involves using the microfluidic ratchet device to sort RBCs based on deformability in order to acquire training data for a deep learning image classification model. The microfluidic ratchet device (**Fig. 1A**) sorts RBCs based on deformability by squeezing cells through a matrix of tapered constrictions (**Fig. 1B**). After microfluidic sorting (**Fig. 1C**), RBCs from each outlet are extracted from the microfluidic device and placed in wells on a 96-well plate (**Fig. 1D**). These wells are then imaged using an optical microscope in brightfield using a 40X objective (**Fig. 1E-F**). The microscopy images are processed and segmented into single cell images (**Fig. 1G**). The resulting datasets are then used to train and test a convolutional neural network for deformability-based image classification (**Fig. 1H**). Finally, the trained deep learning model is used to estimate the deformability of RBCs in a test set. The aggregate RBC deformability is also used to obtain a deep learning-derived deformability profile of the RBC sample (**Fig. 1I**), which is compared to the profile obtained by microfluidic testing (**Fig. 1J**).

**Fig. 1.**
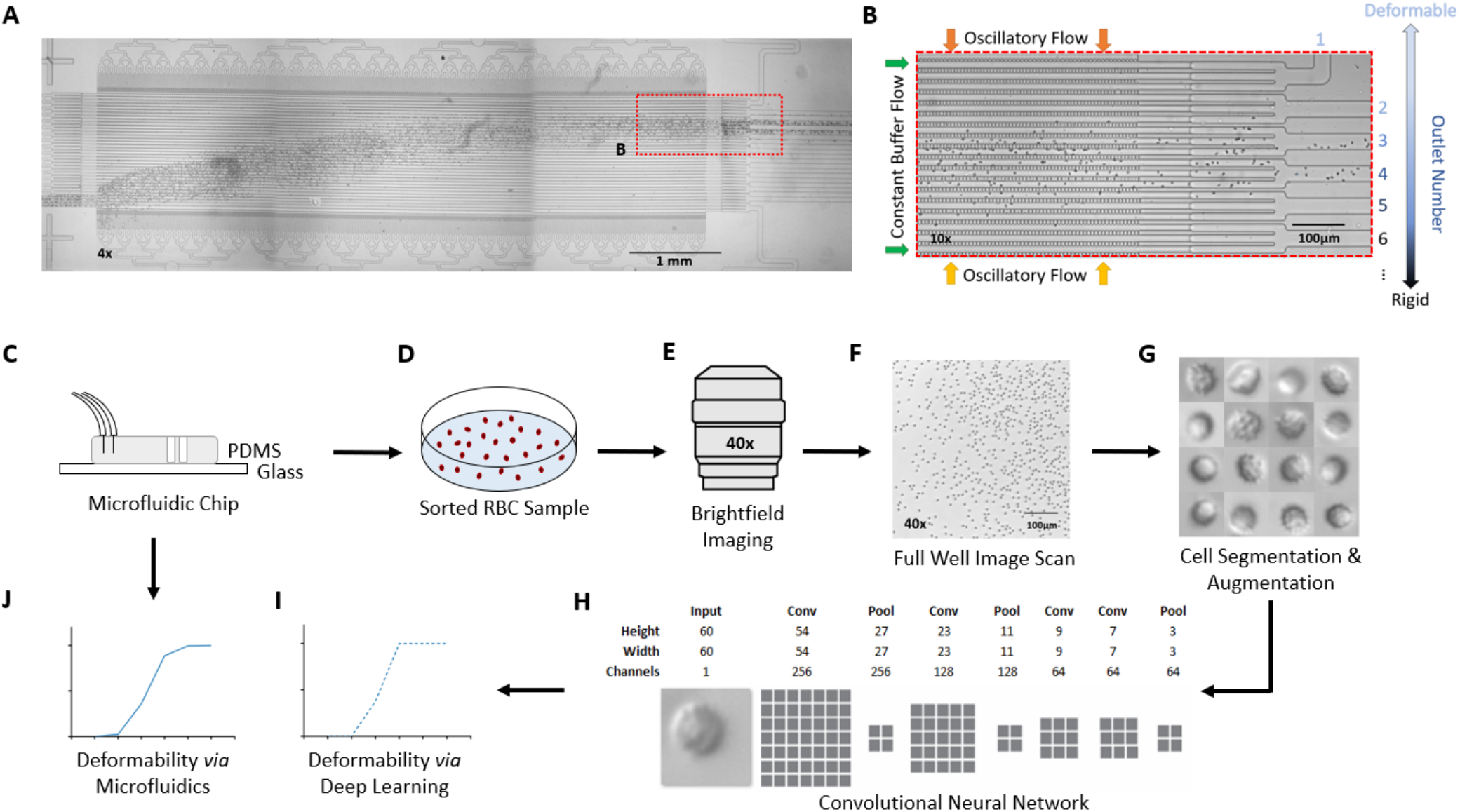
(**A**) Micrograph of the microfluidic ratchet device to sort RBCs based on deformability. (**B**) Closeup of the active region where cells are routed to different outlets. (**C-J**) Overall approach. (**C**) Deformability based sorting using the microfluidic ratchet device. (**D**) Sorted RBC fractions are transferred to a well plate for imaging. (**E**) Brightfield imaging using a 40X objective. (**F**) Example full well image scan. (**G**) Examples of individual segmented RBCs. (**H**) Structure of the convolutional neural network for image-based cell classification. (**I**) RBC Rigidity Score estimated using the CNN. (**J**) Rigidity Score measured by deformability-based cell sorting.

### Deformability based cell sorting

We sorted RBCs based on deformability using the microfluidic ratchet device described previously. ^32–34^ Briefly, RBCs migrate under oscillatory flow through a matrix of micropores with openings ranging from 1.5 µm to 7.5 µm (**Table 1**). The micropores have the same opening in each row, but progressively smaller openings along each column (**Fig. 1A**). When the cells can no longer transit the micropores along a particular row, they instead flow along the row and are directed into one of 12 outlets (**Fig. 1B**). Oscillatory flow within the device dislodges cells that are blocked by the micropores to ensure that the device does not foul. In this way, the RBC sample is fractionated based on deformability at a rate of ∼600 cells per minute.

**Table 1.**
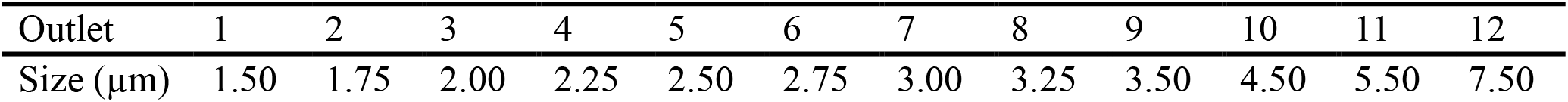
Constriction size for each outlet number.

| Outlet | 1 | 2 | 3 | 4 | 5 | 6 | 7 | 8 | 9 | 10 | 11 | 12 |
| --- | --- | --- | --- | --- | --- | --- | --- | --- | --- | --- | --- | --- |
| Size ($\mu\text{m}$) | 1.50 | 1.75 | 2.00 | 2.25 | 2.50 | 2.75 | 3.00 | 3.25 | 3.50 | 4.50 | 5.50 | 7.50 |

### RBC Rigidity Score (RS)

After deformability-based sorting, the RBC distribution can be shown as a histogram, where a more rightward distribution corresponds to a more rigid RBC sample (**Fig. 2A**). To compare deformability between samples, the RBC distribution can be shown as a cumulative distribution, which allows us to define a Rigidity Score (RS) as the outlet number where the cumulative distribution crosses 50%. Fractional outlet numbers can be determined by linear interpolation between data points greater and less than 50% in the cumulative distribution function. RBC samples from different donors showed significant variability in their RS value. For example, from the ten donors in this study the RS ranged from 2.47 to 3.50 (**Fig. 2B-C**).

**Fig. 2.**
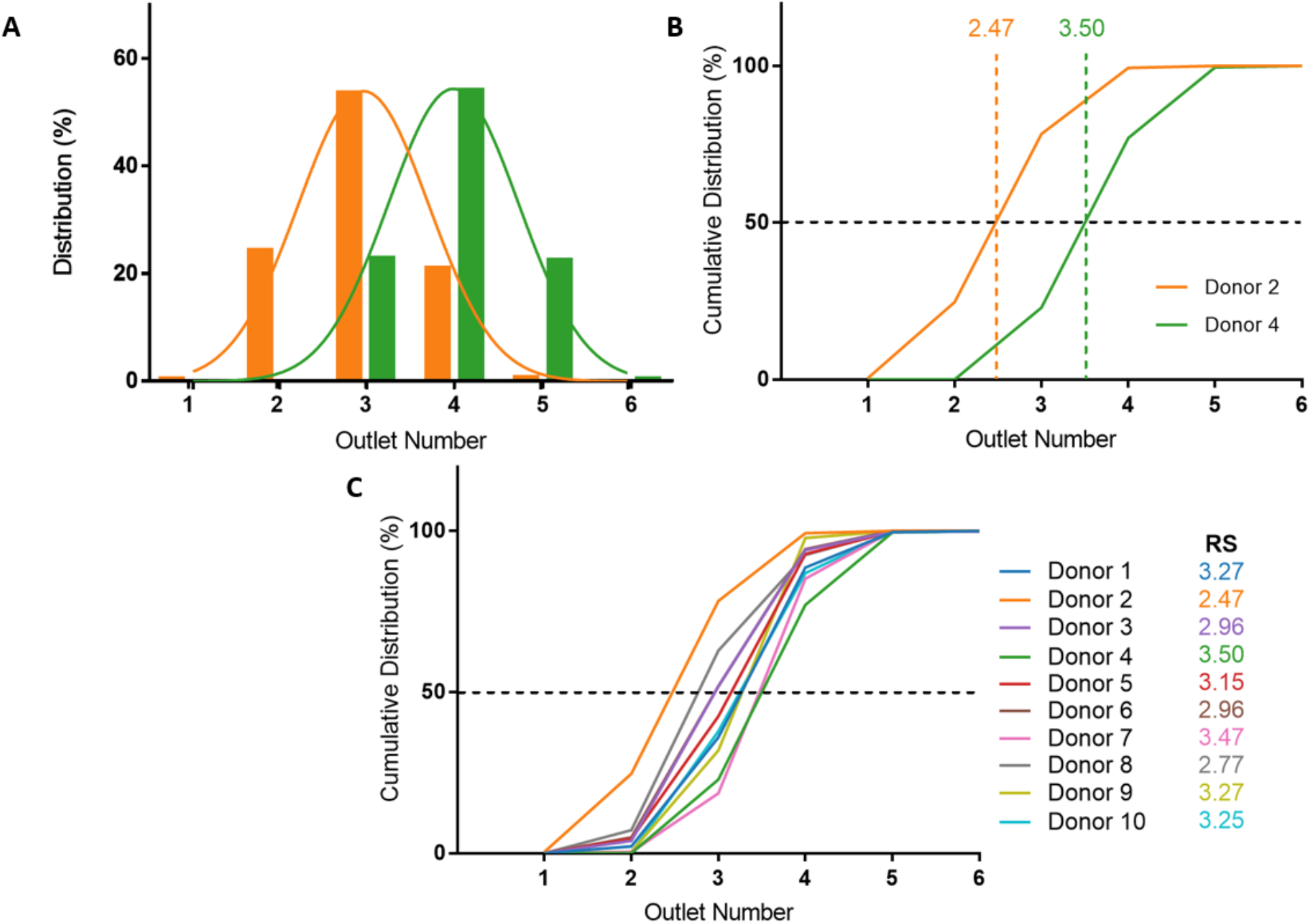
Microfluidic deformability-based sorting results. (**A**) RBC distribution after deformability-based sorting for select donors. Donor 2 is the most deformable sample (orange) and donor 4 is the most rigid sample (green) of the ten donors analyzed. (**B**) Cumulative distribution of RBC deformability from donor 2 and 4. The Rigidity Scores (RS) are measured at the 50% crossover of the cumulative distribution. (**C**) Cumulative distributions and RS for all ten donors.

### Donor-based variability in RBC deformability

Blood donations from ten healthy donors were obtained and RBC samples were sorted based on deformability using the microfluidic ratchet device (**Fig. 2C**). The donors were diverse in terms of blood type and sex (**Table 3**). Seven blood samples were fresh (≤ 3 days after donation), two were in the second week of tube storage, and one was stored in a blood bag for 3 weeks. Donor RBCs were sorted into outlets 1-6, with the majority (>99%) sorted to outlets 2-5. The cumulative distribution of sorted cells to each outlet is presented in **Fig. 2C**. These deformability curves and RS are donor-specific and can be reliably measured in repeated experiments using replicate microfluidic devices. ^9^

### Optical microscopy imaging for deep learning

After deformability-based cell sorting, the sorted cells are extracted from the microfluidic device by pipetting and placed in 96-well imaging plate. Samples from each outlet were split in half and placed in two wells to introduce additional variance in the imaging conditions. These variations include a greater variety of light conditions based on location of cells in the well, cells with different thicknesses of suspension fluid due to its meniscus, and different imaging conditions resulting from differences in focus and exposure time. Full image scans were conducted on each well using a 40X objective and a DS-Qi2 camera on a Nikon Ti-2E inverted microscope, capturing brightfield images of 2424×2424 pixels (**Fig. 1E-F**). Image captures near the edge of the wells were often out of focus and were discarded prior to segmentation.

### Segmentation

To perform deep learning classification of individual RBCs, we developed a Python program to extract 60×60 pixel image patches each containing a single RBC (**Fig. 1G**). Cells are located using a Sobel operator for edge detection and Otsu multi-thresholding. A centre of mass measurement is conducted to centre the identified cell for image cropping. After the cells are identified and cropped, they proceed through a selection algorithm that keeps images with a single cell centred in the image and rejects images with multiple cells. Further, the resulting selected cropped images are manually audited and any remaining images with multiple cells or those out of focus are removed. This segmentation procedure resulted in datasets containing 20,000 to 70,000 single cell images for each donor (**Table 2**).

**Table 2.**
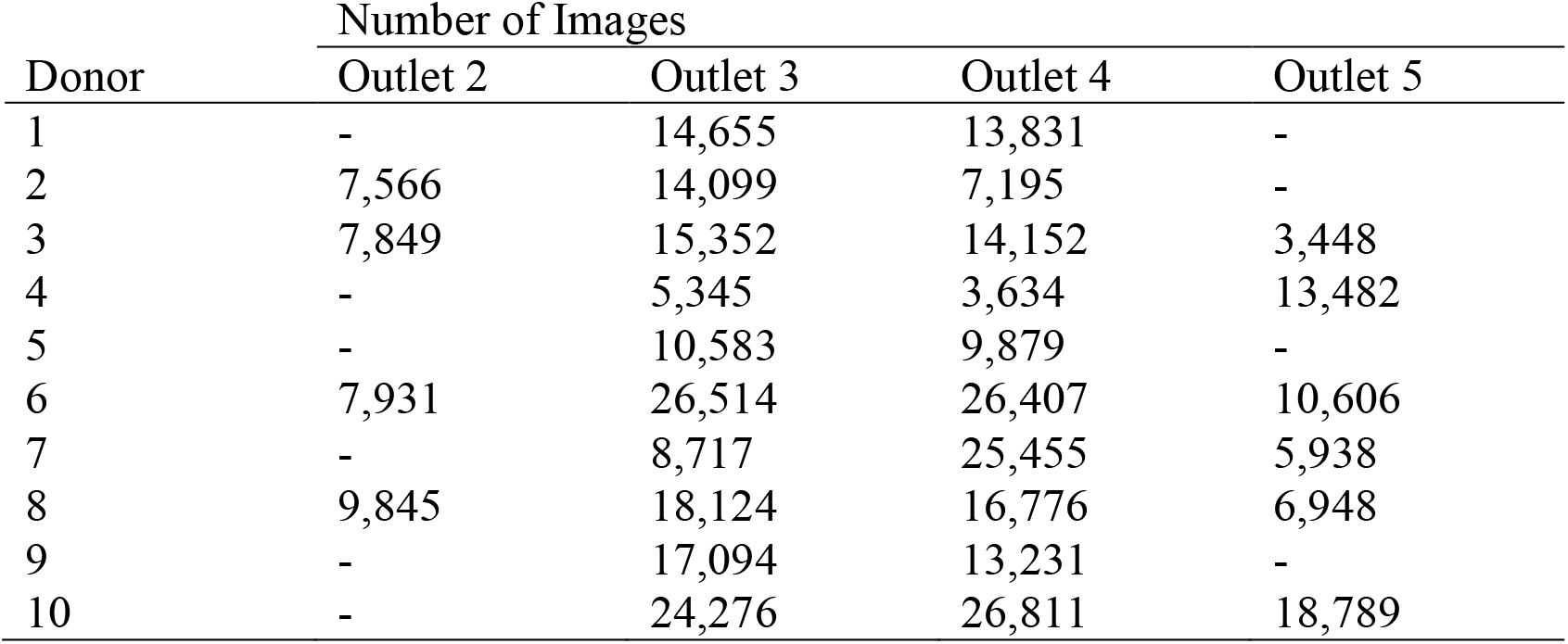
Number of unique segmented single cell images in each outlet for all ten donors. From these datasets, *deformable* (outlets 2 and 3) and *rigid* (outlets 4 and 5) images are split for training and testing, and are subsequently augmented for class balancing.

### Network design

We designed a convolutional neural network (CNN) to conduct image feature extraction and classification using the Keras library in TensorFlow (**Fig. 1H**). For feature extraction, the model utilizes a series of 4 convolutional layers and 3 max pooling layers. The initial convolutional layer kernel size is 7×7, the second is 5×5, and the final two are 3×3. Each convolutional layer is followed by batch normalization and ReLU activation. The latter classification section consists of 3 fully connected layers and a final smaller fully connected output layer. The 3 fully connected layers are followed by batch normalization, ReLU activation, and 20% dropout. The output layer uses a SoftMax error function for backpropagation during training. The network utilizes a binary cross-entropy loss function and stochastic gradient descent for optimization.

### Training and validation

The CNN was trained using single-cell images with true deformability labels determined by microfluidic deformability-based cell sorting. The CNN utilized balanced training classes of 10,000 images per class per donor. Cell images were augmented by a random integer multiple of 90-degree rotation to capture different cell orientations and lighting characteristics (**Fig. 1G**). Classes with fewer than 10,000 images were up-sampled, and classes with greater than 10,000 images were sub-sampled. The model was trained for a minimum of 30 epochs for Donor 1 and up to 80 epochs for Donor 7. There was significant variation in the model’s ability to converge during training between different donors, illustrated by the variation in training epochs. The final training accuracies for each donor is shown in **Table 3** and **Fig. 3D**. The training results were validated using a held-out validation split of 10% of the training set. Using the validation dataset, the model was evaluated using the corresponding validation convergence after every epoch for hyperparameter tuning.

**Table 3.** Donor characteristics, deep learning results, and comparison of microfluidic and deep learning determined Rigidity Scores (RS).

| Donor | Blood Type | Sex | Outlets | Training Accuracy (%) | Validation Accuracy (%) | Testing Accuracy (%) | Microfluidic RS | Deep Learning RS |
| --- | --- | --- | --- | --- | --- | --- | --- | --- |
| 1 | A- | F | 3, 4 | 92 | 92 | 92 | 3.27 | 3.20 |
| 2 | A+ | M | 2, 3, 4 | 96 | 96 | 95 | 2.47 | 2.67 |
| 3 | O- | M | 2, 3, 4, 5 | 86 | 86 | 86 | 2.96 | 3.09 |
| 4 | A+ | M | 3, 4, 5 | 99 | 99 | 98 | 3.50 | 3.34 |
| 5 | B+ | F | 3, 4 | 85 | 83 | 82 | 3.15 | 3.17 |
| 6 | B+ | M | 2, 3, 4, 5 | 78 | 66 | 68 | 2.96 | 3.07 |
| 7 | B+ | M | 3, 4, 5 | 88 | 76 | 71 | 3.47 | 3.20 |
| 8 | B+ | F | 2, 3, 4, 5 | 83 | 87 | 72 | 2.77 | 2.91 |
| 9 | O* | F | 3, 4 | 97 | 97 | 95 | 3.27 | 3.22 |
| 10 | B+ | F | 3, 4, 5 | 91 | 82 | 82 | 3.25 | 3.06 |

\* Donor Rhesus factor unknown

**Fig. 3.**
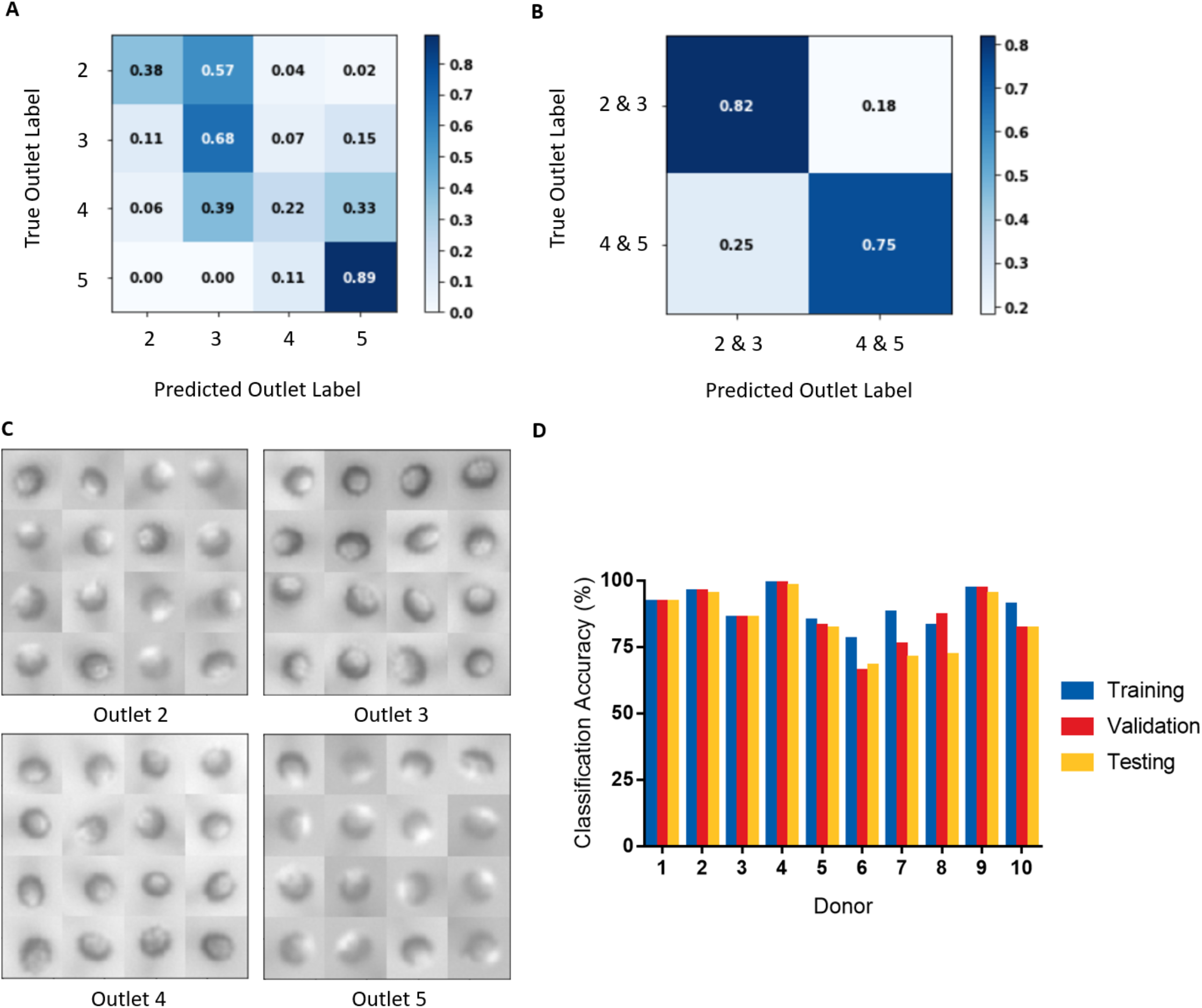
(**A**) Normalized confusion matrix for donor 3 when RBCs from each outlet were considered as a separate class. (**B**) Normalized confusion matrix for donor 3 when RBCs from outlets 2 and 3 (*deformable*) were pooled into a single class, and RBCs from outlets 4 and 5 (*rigid*) were pooled into a single class. Classification accuracy is greatly increased with outlets pooled. (**C**) Example images from outlets 2-5 for donor 3. (**D**) Image classification training, validation, and testing accuracies for all donors.

### Classification

We initially used our CNN to classify cells from microscopy images based on the outlet they were sorted to. However, classifying cells in this manner resulted in poor classification accuracies (**Fig. 3A**). This result likely derives from there being substantially fewer single cell images from outlets 2 and 5, compared to outlets 3 and 4 (**Table 2**). To create balanced classes of 10,000 images, substantial up-sampling was performed on cells from outlets 2 and 5, requiring many repeated cells. As a result, the variety of cell images seen by the model for these classes were significantly limited. Interestingly, more misclassification occurred between adjacent outlets (**Fig. 3A**), indicating that the model was learning some common deformability-based cell features.

To improve our classification accuracy, we collapsed the image data from outlets 2 and 3 together and outlets 4 and 5 together to create classes of *deformable* and *rigid* cells. By collapsing the classes in this manner, the datasets are more robust as additional cell images are available for augmentation, which required less up-sampling. This binary classification method resulted in substantially improved deformability image predictions (**Fig. 3B**). An additional advantage of this binary classification method is that inter-donor comparisons are more appropriate as all donors have cells sorted to outlets 3 and 4, but not all have cells sorted to outlets 2 or 5. Interestingly, since differences in deformability is not a feature that can be easily decerned by a human observer, a large and robust dataset is required for training and testing (**Fig. 3C**). Using this classification scheme, training accuracies ranged 78-99% and final validation accuracies ranged 66-99%, shown in **Table 3** and **Fig. 3D**.

### Testing

The testing datasets are comprised of 20% of the overall segmented data and were separated from the training and validation sets prior to augmentation. This process ensured that there are no cell image repeats between the training and testing sets. For each donor, the images were augmented using the same method used in training and were up-sampled or down-sampled to obtain balanced testing sets of 2,000 images per class. Among our donor population, we observed testing accuracies ranging 68-98% with an aggregate mean (± SD) of 84 ± 11% (**Table 3** and **Fig. 3D**). For each donor, we observed that testing accuracy corresponded well with validation accuracy.

### Saliency maps

A key consideration for image classification using deep learning is whether classification was driven by imaging artifacts, such as lighting, sample preparation, image acquisition parameters, and position in the imaging well. ^55^ To resolve this potential issue, we generated saliency maps to assess if the model learned relevant cell morphological features. A saliency map is a visual representation of the spatial support of a particular class to indicate which pixels in a given image had the greatest effect on the classification probability. ^56^ As shown in **Fig. 4**, the saliency map shows that pixels having the most influence on classification corresponded to those clustered around the cell itself, rather than surrounding regions. This result confirms that our model is classifying RBC deformability based on cell morphological features.

**Fig. 4.**
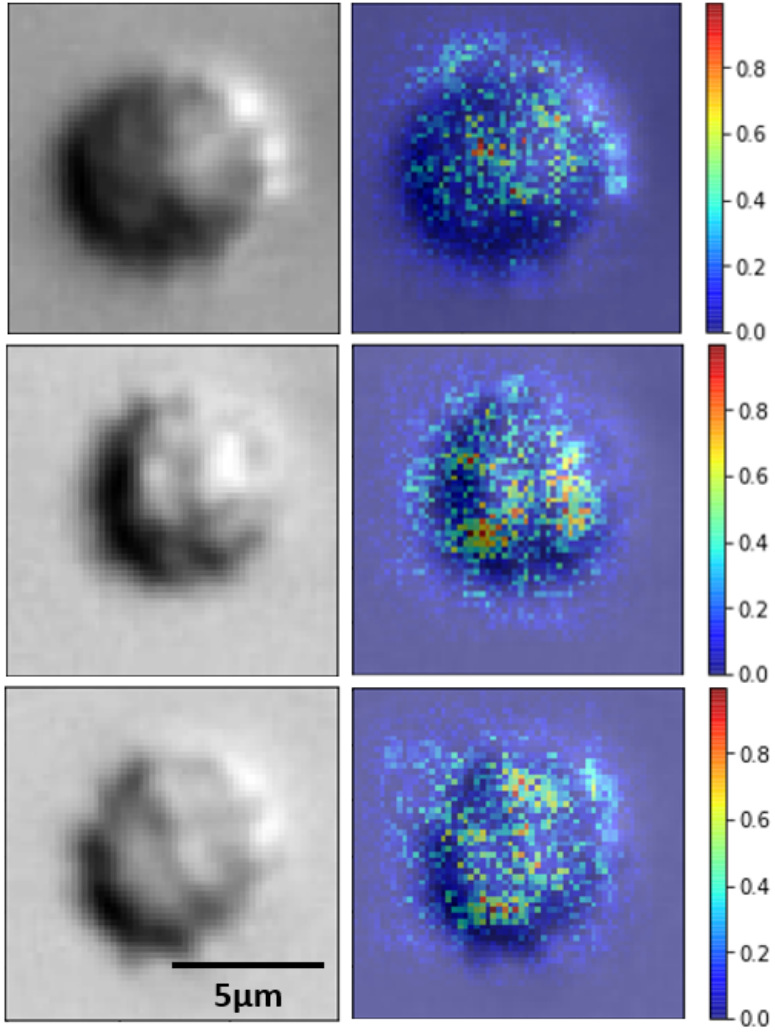
Original images (left) and saliency maps (right) of randomly sampled RBCs from donor 1. The saliency map indicates the strength of pixel contributions to the final classification output by the CNN. Warmer colours indicate greater contribution and cooler colours indicate lesser contribution.

### Using deep learning to determine Rigidity Scores of RBC samples

By classifying RBCs into *deformable* and *rigid* fractions, we can use this result to estimate the RS for each RBC sample. RBCs classified by deep learning as *deformable* and *rigid* classes are assigned to outlet 3 and 4, respectively. This scheme is justified for the ten donors studied here since the vast majority (86%) of cells from all these donors were sorted into outlets 3 and 4, with the reminder sorted into outlets 2 and 5. This approach also kept the RS calculations consistent between all donors as not every donor had cells sorted to outlets 2 or 5, but all donors had cells sorted to outlets 3 and 4. After classification, the RS is then determined as before by linearly interpolating the cumulative distribution deformability curve to find the outlet number at the 50% crossover frequency.

Comparing the cumulative distributions and RS obtained by cell sorting using the microfluidic ratchet device with the RS estimated by deep learn showed a strong agreement (**Fig. 5**). Specifically, the measured and estimated RS values deviated between a minimum 0.02 (donor 5) to a maximum 0.27 (donor 7) with a mean of 0.13 ± 0.08. Expressed differently, the deep learning estimated RS deviated from the microfluidic RS across the ten donors by a mean of 4.4 ± 2.5%. Previous work with this microfluidic device has shown a standard deviation for RS of 0.17 (6.5%) across five different tests on the same sample, ^9^ indicating the level of deviation seen here is within acceptable variation in RS resulting from random sampling and manufacturing. Furthermore, we plotted the RS acquired by deep learning against RS acquired by microfluidics for the ten donors and found a high degree of correlation between the two methods (**Fig. 5K**), with a Pearson’s correlation of *r* = 0.93 and *p* < 0.0001.

**Fig. 5.**
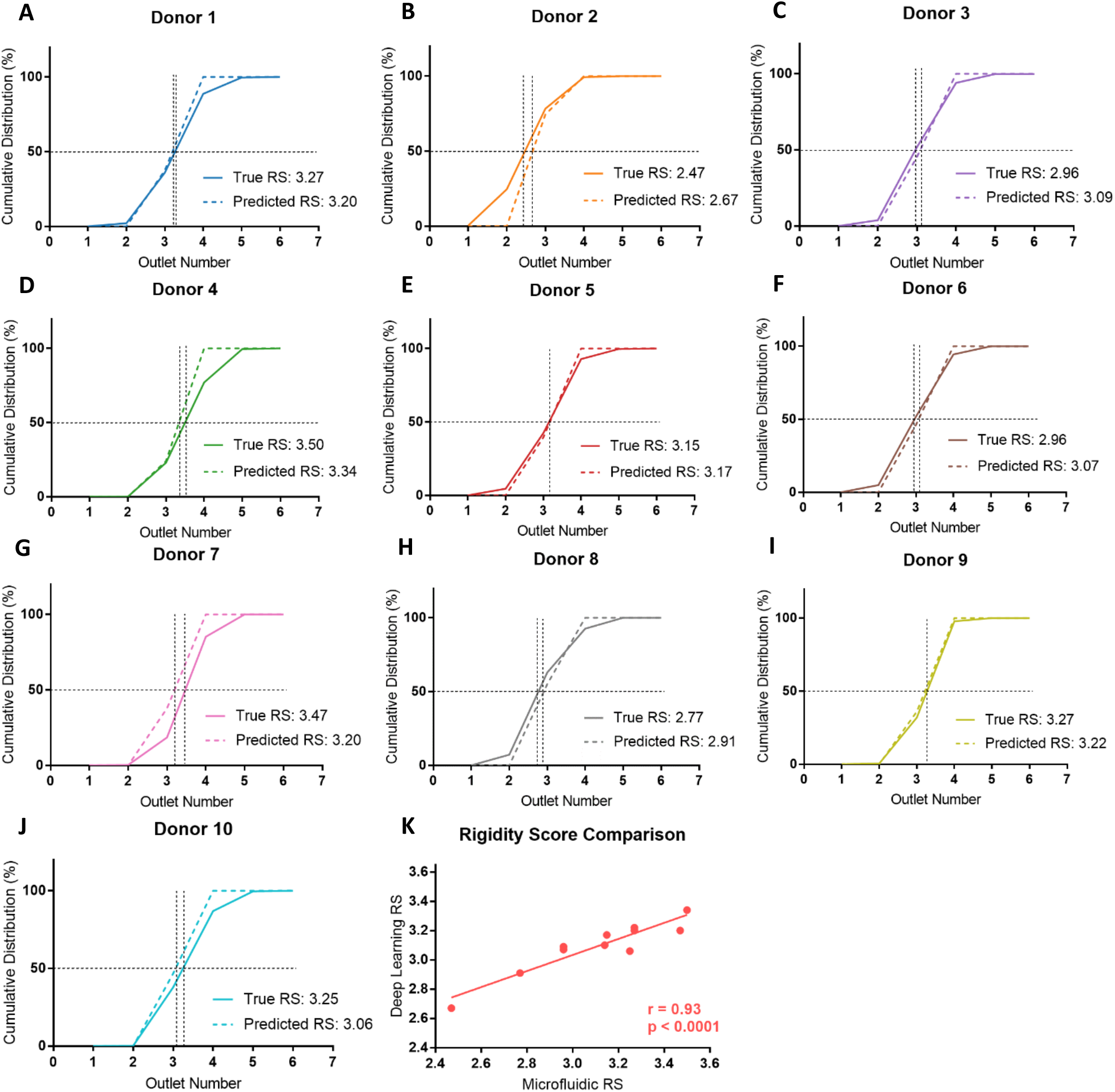
(**A-J**) Comparison of microfluidic (solid lines) and deep learning (dashed lines) derived RBC deformability cumulative distributions and RS for all ten donors. (**K**) Relationship between Rigidity Scores determined by microfluidics and deep learning methods for all ten donors. The deep learning RS values are strongly correlated to the microfluidic RS values (*r* = 0.93) and this relationship is statistically significant (*p* < 0.0001).

To assess whether cell sorting using the microfluidic ratchet device may have altered the RBCs, we used the donor-specific trained CNN to classify unsorted cells from donor 2, 3, and 4. This test also assessed the model’s generalizability by applying a testing dataset acquired in a separate experimental procedure from the training data. Donor 2, 3, and 4 were selected for assessment as these donors represent the full range of donor RS: donor 2 is most deformable (RS = 2.47), donor 3 is in the middle (RS = 2.96) and donor 4 is the most rigid (RS = 3.50). As before, the cumulative distribution and RS obtained by cell sorting and deep learning were similar with the difference in RS for donor 2, 3, and 4 being 0.04 (1.6%), 0.26 (8.8%), and 0.09 (2.6%), respectively (**Fig. 6**). In summary, these results show that our deep learning model is a robust and generalizable for classifying the deformability of RBCs acquired and processed separately from the training dataset.

**Fig. 6.**
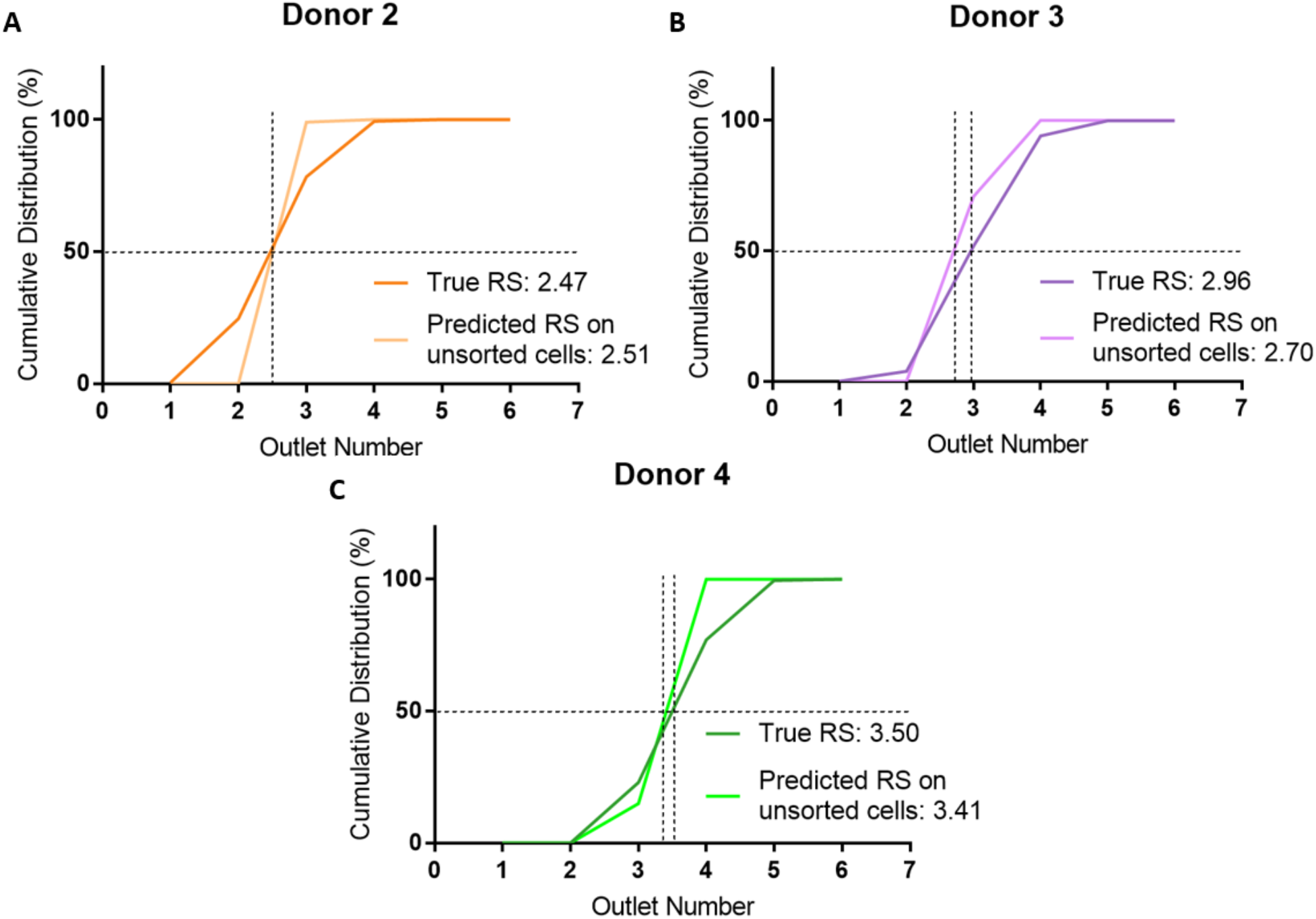
Generalization of the deep learning network applied to unsorted cell images (light lines) compared to microfluidic-derived deformability cumulative distributions (dark lines) for donor 2 (**A**), 3 (**B**), and 4 (**C**). The model generalizes well on the unsorted datasets, illustrated by similar cumulative distribution plots and Rigidity Scores between the ground truth microfluidic results and the deep learning model tested on an unsorted RBC sample.

## DISCUSSION

This study developed a deep learning method to infer RBC deformability that bypasses the need for complex and time-consuming physical measurement. By sorting RBCs into fractions based on deformability and imaging cells from each fraction, we were able to use deep learning to classify RBCs based on deformability with a mean testing accuracy of 84%. Furthermore, we used this approach to estimate RBC Rigidity Scores (RS), which deviated only by a mean of 4.4% from physical measurement using the microfluidic ratchet device. We previously showed significant inter-donor variability in RBC deformability, ^9^ which were successfully captured by our deep learning model. To ensure our inference of RBC deformability was robust to changes in well location, lighting conditions, and presence of debris during imaging, we introduced additional data variance by purposefully dividing RBC specimens into multiple imaging wells and augmenting cell images by rotation during database creation. We further used saliency maps to confirm that cellular features, not imaging artifacts, were learned for RBC classification. Finally, the generalizability of this method was investigated by using a donor-specific trained model to evaluate the deep learning derived deformability profile from unsorted donor RBCs. The RS obtained by this approach showed strong agreement with microfluidic sorting, deviating by 1.6-8.8% from the microfluidic measurement, which is comparable to previously reported variability of microfluidic ratchet measurement (6.5% from five independent measurements). ^9^

Measuring RBC deformability using microscopy imaging and deep learning has several important advantages over physical measurement. First, microfluidic devices used to measure RBC deformability have limited throughput owing to the need to flow single cells within the device. In contrast, using machine learning to infer RBC deformability allows for evaluation of cells densely seeded within imaging wells (∼1,500 cells imaged per minute). Second, a major barrier to physical measurement of RBC deformability is that these methods are prohibitively difficult, time consuming, and require specialized equipment. ^35,57,58^ In contrast, microscope systems are ubiquitous in both research and clinical laboratories. Finally, while there are several methods available for physical measurement of RBC deformability, there are no accepted standards by which to compare studies. The universality of microscope systems could offer an approach to standardize RBC deformability measurements in order to extend studies across multiple centers. In summary, RBC assessment by machine learning is more accessible, simpler to perform, and more standardizable compared to physical RBC deformability measurements.

To our knowledge, this work describes the first instance of employing deep learning to measure RBC deformability. Deep learning has been used previously to characterize other cellular properties of RBCs. For example, Doan et al. ^38^ trained a deep learning model to classify unlabeled images of stored RBCs into seven morpho-types with 76.7% accuracy, which was comparable to 82.5% agreement in manual classification by experts. Other studies trained deep learning models to identify RBCs from patients with malaria, ^39–44^ sickle cell disease, ^45–50^ and thalassemia, ^51–53^ based on visually identifiable changes in RBC morphology. Our application of machine learning in RBC deformability measurement deviates from these previous efforts because cellular features corresponding to deformability are beyond human perception. This result expands on our previous study using deep learning to distinguish between cell lines that lack readily differentiable features to a human observer, ^54^ further supporting our belief that imperceivable cellular parameters, such as changes in biophysical or metabolic cell state, may be measurable using deep learning. Potential future applications of this work include assessment of RBC units prior to transfusion in order to preferentially allocate stored blood bags to the most appropriate recipients.

## METHODS

### RBC sample collection and preparation

This study was approved by the University of British Columbia Clinical Research Ethics Board (UBC REB# H19-01121) and the Canadian Blood Services Research Ethics Board (CBS REB# 2019-029). Donors self-identified as healthy and between ages 18-70 provided fresh RBCs in citrate tubes (n=3) or blood bags (n=7). Donors were diverse in terms of blood type and sex (**Table 3**). Blood sample components were separated by centrifuging at 3900 rpm for 8 minutes at room temperature. Plasma supernatant and leukocyte buffy coat were removed and disposed. The RBC pellet was resuspended and washed three times using Hanks balanced salt solution (HBSS, Gibco) with 0.2% Pluronic solution (F127, MilliporeSigma) by centrifuging at 1800 rpm for 5 minutes. After all supernatant and leukocytes are removed, the RBC pellet was diluted to 1% hematocrit in HBSS + 0.2% Pluronic for infusion into the microfluidic device.

### Microfluidic ratchet device manufacture

The manufacture of the microfluidic devices has been described previously. ^59,60^ The master device mold was created using photolithographic microfabrication and was used to create a secondary master polyurethane mold fabricated out of Smooth-Cast urethane resin (Smooth-Cast ONYX SLOW, Smooth-On) as described here. ^61^ Single-use microfluidic ratchet devices were molded from the secondary master using PDMS silicone (Sylgard-184, Ellsworth Adhesives) mixed at a 10:1 ratio with the PDMS curing agent (Sylgard-184, Ellsworth Adhesives). The PDMS molded devices were then cured for two hours at 65°C. The cured PDMS devices were removed from the molds and manually punched with 0.5 and 3.0 mm hole punches (Technical Innovations). A thin PDMS silicone (RTV 615, Momentive Performance Materials LLC) layer was manufactured to seal the device’s microstructures. This layer was produced by spin coating uncured PDMS on a 100 mm silicon wafer at 1500 rpm for 1 minute, then was cured for 2 hours at 65°C. The Sylgard-184 PDMS microstructure mold was bonded to the RTV 615 thin PDMS layer using air plasma (Model PDC-001, Harrick Plasma). Finally, the composite sealed microstructure mold was then bonded to a 75×50 mm glass slide (Corning) using air plasma.

### Microfluidic device operation

The microfluidic ratchet sorting device is operated via 4 pressurized fluidic inputs. A horizontal crossflow moves the sample towards the outlets, while a vertical (relative to **Fig. 1A-B**) oscillating pressure system squeezes cells through the tapered constrictions and de-clogs others unable to pass through. Before the RBC sample is infused, the device is buffered with HBSS with 0.2% Pluronic-F127 solution through the horizontal crossflow inlet at high pressure (300 mbar) for 15 minutes. Once the device is buffered, 10µL of HBSS with 0.2% Pluronic-F127 solution is pipetted into each outlet (which are open to atmospheric pressure) to improve ease of removal of each RBC deformability sample after sorting. The RBC sample for each donor is suspended at 1% hematocrit in HBSS with 0.2% Pluronic-F127 and then infused into the microfluidic device at 40-45 mbar through the sample inlet. The sample flows through the constriction matrix via the horizontal crossflow pressure (55-60 mbar) and oscillatory pressure of 175 mbar upwards and 162 mbar downwards (relative to **Fig. 1A-B**). The oscillation cycle of these pressures occurs over 5 seconds: 4 seconds of upward pressure flow for filtration, then 1 second of downward flow for declogging. The sorting process throughput is approximately 600 cells per minute; the device is run for 60-90 minutes, resulting in over 30,000 sorted cells. After the cells are sorted through the constriction matrix, the cells proceed to one of 12 distinct deformability outlets. The distribution of sorted cells is determined by capturing images as the cells exit the constriction matrix and counting the cells manually using ImageJ. ^62^ The distribution can also be determined by video analysis of the cells travelling through the constriction matrix exit channels towards the outlets, or by removing the cells in the outlets by pipetting for counting. Sorted RBCs suspended in 10 µL of HBSS with 0.2% Pluronic-F127 from each outlet are removed by pipetting and placed in a 96 well plate (VWR International, LLC) for imaging.

### Microbead sorting validation

Microbead sorting validation was conducted to ensure device manufacturing and sorting consistency between different devices and users. To mimic deformable RBCs, 1.53 µm polystyrene beads (Cat #17133, Polysciences Inc.) were infused into the microfluidic device at 0.1% concentration in HBSS with 0.2% Pluronic F127 and 0.2% TWEEN-20 (MilliporeSigma) to prevent bead aggregation. The bead solution was run through the microfluidic device for 20 minutes, images were captured as the beads exited the matrix sorting region, and the distribution was determined by manually counting beads in ImageJ. Users #1 and #2 conducted 14 total tests using devices from five different master molds. Intra- and inter-user microbead sorting was consistent (**Fig. S1**).

### Image acquisition

After microfluidic sorting, sorted RBCs were removed from the microfluidic device and transferred to a 96-well flat-bottom plate (VWR International, LLC). Samples of sorted cells from each outlet were divided evenly and placed into two separate wells to provide cell images captured with variations in well location distribution and automatically determined imaging parameters (e.g. auto-exposure and auto-focus) to produce robust datasets. Full image scans of each well in 40X brightfield were acquired using a Nikon Ti-2E inverted microscope and NIS Elements software.

Illumination for the brightfield images was implemented by using the built-in Ti-2E LED. Gain, exposure, and vertical offset were automatically determined by built-in NIS Elements functions for consistency and to avoid user bias. Components of the full image scan were 2424×2424 pixel BMP images with 24-bit depth.

### Segmentation and augmentation

Each full scan image was segmented using a custom computer vision segmentation algorithm. Individual cells were identified using a watershed algorithm and were segmented into 60×60 pixel PNG images with 8-bit depth. Segmented images with multiple or partial cells were manually removed. Resultant single cell images from each donor were split at an 80:20 ratio per class to create separate training and testing datasets. After database splitting, images were augmented by a random multiple of 90° rotation (0°, 90°, 180°, or 270°). This augmentation allowed for the building of balanced training (10,000 images per outlet) and testing (2,000 images per outlet) datasets for each donor. In addition, different lighting conditions were observed depending on the location of the cell in the well. By augmenting the cells by rotation this potential data confounder is mitigated.

### CNN model

A convolutional neural network, shown in **Fig. 1H**, was designed in Python using the Keras library in TensorFlow. The network accepts a 1-channel input of 60×60 pixels. The model begins with a 256-channel convolution layer with a kernel size of 7 and a stride of 1. Next, the model uses a 2×2 max-pooling layer with a stride of 2. The next layer is a 128-channel convolutional layer with a kernel size of 5×5 and a stride of 1, followed by another max-pooling layer of size 2×2 with a stride of 2. Next are two 64-channel convolutional layers in series, each with kernel sizes of 3 and strides of 1. These layers were followed by a max-pooling layer of size 2×2 with a stride of 2. Then, the layer outputs were flattened into a 1-dimensional array for connection to the fully connected layers. Each of the four convolutional layers were followed by ReLU activation and batch normalization. Then, three 128-node fully connected dense layers, consisting of ReLU activation and 20% dropout, were used by the model to learn on the identified features from the earlier convolutional layers. The network outputs two nodes, one per class, with a softmax (normalized exponential) error function for backpropagation.

### Training environment

The segmentation and deep learning software were run on a Lenovo Thinkpad X1 Extreme operating Windows 10 Pro with an Intel® Core™ i7-8850H CPU running at 2.60 GHz. The computer used 32.0 GB DDR4 RAM running at 2933 MHz. The graphics card used was a 4096 MB GeForce GTX 1050 Ti with Max-Q Design. Training and testing were conducted in Python 3.6.12 utilizing the TensorFlow 2.1.0 library.

### Training

For each execution of the network, training occurred for 30 to 80 epochs with stochastic gradient descent optimization and a learning rate between 0.1 and 0.0001. The appropriate number of epochs and learning rate were determined iteratively to find the best combination for training convergence and validation accuracy for each donor/dataset combination. Training concluded after there was no improvement in the loss after the past five epochs. A batch size of 32 and a categorical cross entropy loss function from the Tensorflow Keras library (version 2.2.4) were used. The error function used for backpropagation was the SoftMax function. 10% of the training data was split for validation during training.

### Testing

To verify the accuracy on the 10% of the training dataset that was held-out for validation, testing occurred on the 2,000 images per outlet from the separate testing dataset. This dataset was split from the training set prior to augmentation, ensuring images used for testing were previously unseen by the network. In addition to deep learning testing conducted on microfluidic sorted cells, unsorted cell images from donor 2 (6,307 images), 3 (9,003 images), and 4 (2,562 images) were also assessed.

## Acknowledgments

We are grateful to Canadian Blood Services’ blood donors who made this research possible.

